# Functional signatures of evolutionarily young CTCF binding sites

**DOI:** 10.1101/2020.01.31.928119

**Authors:** Dhoyazan Azazi, Jonathan M. Mudge, Duncan T. Odom, Paul Flicek

## Abstract

The introduction of novel CTCF binding sites in gene regulatory regions in the rodent lineage is partly the effect of transposable element expansion. The exact mechanism and functional impact of evolutionarily novel CTCF binding sites are not yet fully understood. We investigated the impact of novel species-specific CTCF binding sites in two *Mus* genus subspecies, *Mus musculus domesticus* and *Mus musculus castaneus,* that diverged 0.5 million years ago. The activity of the B2-B4 family of transposable elements independently in both lineages leads to the proliferation of novel CTCF binding sites. A subset of evolutionarily young sites may harbour transcriptional functionality, as evidenced by the stability of their binding across multiple tissues in *M. musculus domesticus* (BL6), while overall the distance of species-specific CTCF binding to the nearest transcription start sites and/or topologically-associated domains (TADs) is largely similar to *musculus*-common CTCF sites. Remarkably, we discovered a recurrent regulatory architecture consisting of a CTCF binding site and an interferon gene that appears to have been tandemly duplicated to create a 15-gene cluster on chromosome 4, thus forming a novel BL6 specific immune locus, in which CTCF may play a regulatory role. Our results demonstrate that thousands of CTCF binding sites show multiple functional signatures rapidly after incorporation into the genome.

## INTRODUCTION

Genetic differences within and between species predominantly lie in the noncoding sequence of the regulatory regions of the genome, whose function and significance remain poorly understood (1–3). While interspecies comparisons of mammalian genomes have revealed that protein-coding genes have been subject to strong selective pressures (4), tissue-specific transcription factor binding diverges more frequently between species (5–8).

CCCTC-binding factor (CTCF) is a ubiquitously expressed 11 zinc-finger master genome organizer (9) shared between all vertebrates (10). It plays a part in many basic cellular roles including transcriptional activation and repression (11, 12), X-inactivation (13), establishing 3D genome architecture (14), enhancer insulation (15), and alternative splicing (16). The importance of these functions is illustrated by CTCF knockout being embryonic lethal (17), and tissue-specific conditional knockouts having dramatic developmental abnormalities (18, 19). The CTCF protein itself shows a remarkable degree of evolutionary conservation, with 93% amino acid identity of the full protein sequence between human and chicken and 100% identity in its DNA binding domain (20, 21).

One of the most important functions of CTCF is to help establish 3D genome structure through interaction with the cohesin complex (22–25). The colocalised binding of CTCF and the cohesin complex can create chromatin loops, demarcated by two CTCF molecules bound to the genome and stabilised by cohesin (26–28). This gives rise to topologically associating domains (TADs) (13), demarcated by CTCF sites deeply conserved across mammals (29), and with mostly invariant positions across species and tissues (30, 31).

Many changes in the regulatory non-coding genome between species are the consequences of the co-option of repetitive sequences for active binding of transcription factors (32–38). Across mammals, the evolution of CTCF binding has been driven by repeated waves of expansions of transposable elements that deposited its binding motif in novel genomic locations (35–37). Specifically within the mouse lineage, CTCF binding motifs have recently been spread through the short nuclear interspersed elements (SINEs) family of transposable elements. Similar repeat-driven transcription factor binding site birth expansions have been observed for other tissue-specific transcription factors in stem cells (34) and in pregnancy associated tissues (39), suggesting that repeat expansions are a common mechanism used to remodel mammalian genomes (32). However, the potential functional roles of very young insertions of transcription factor binding sites via repeats, their function and genomic characteristics have not yet been well characterised.

Leveraging the availability of high-quality genome sequences from laboratory strains and species within the *Mus* genus created by the Mouse Genomes Project (40–43), we illustrate how repetitive elements have remodelled CTCF binding in two *Mus* genus subspecies sharing a common ancestor 0.5 million years ago (MYA): *Mus musculus domesticus* (C57BL/6J or BL6) and *Mus musculus castaneus* (CAST) (Figure 1A). We found that almost half of the subspecies-specific CTCF binding sites are from repeat origin but have signatures of function and genomic occupancy patterns that are largely similar to older CTCF sites common between the subspecies. We next identified a subset of these subspecies-specific sites that was bound in multiple tissues and exhibit heightened recruitment of cohesin-complex subunits, suggesting active participation in loop formation and higher functionality of these sites. We also found a cluster of interferon genes with BL6 subspecies-specific CTCF and cohesin colocalised binding sites on mouse chromosome 4 that apparently arose via a recent tandem duplication event. Taken together, these results demonstrate the pace at which evolutionarily young CTCF binding sites appear in the genome and acquire functional signatures.

**Figure 1.**
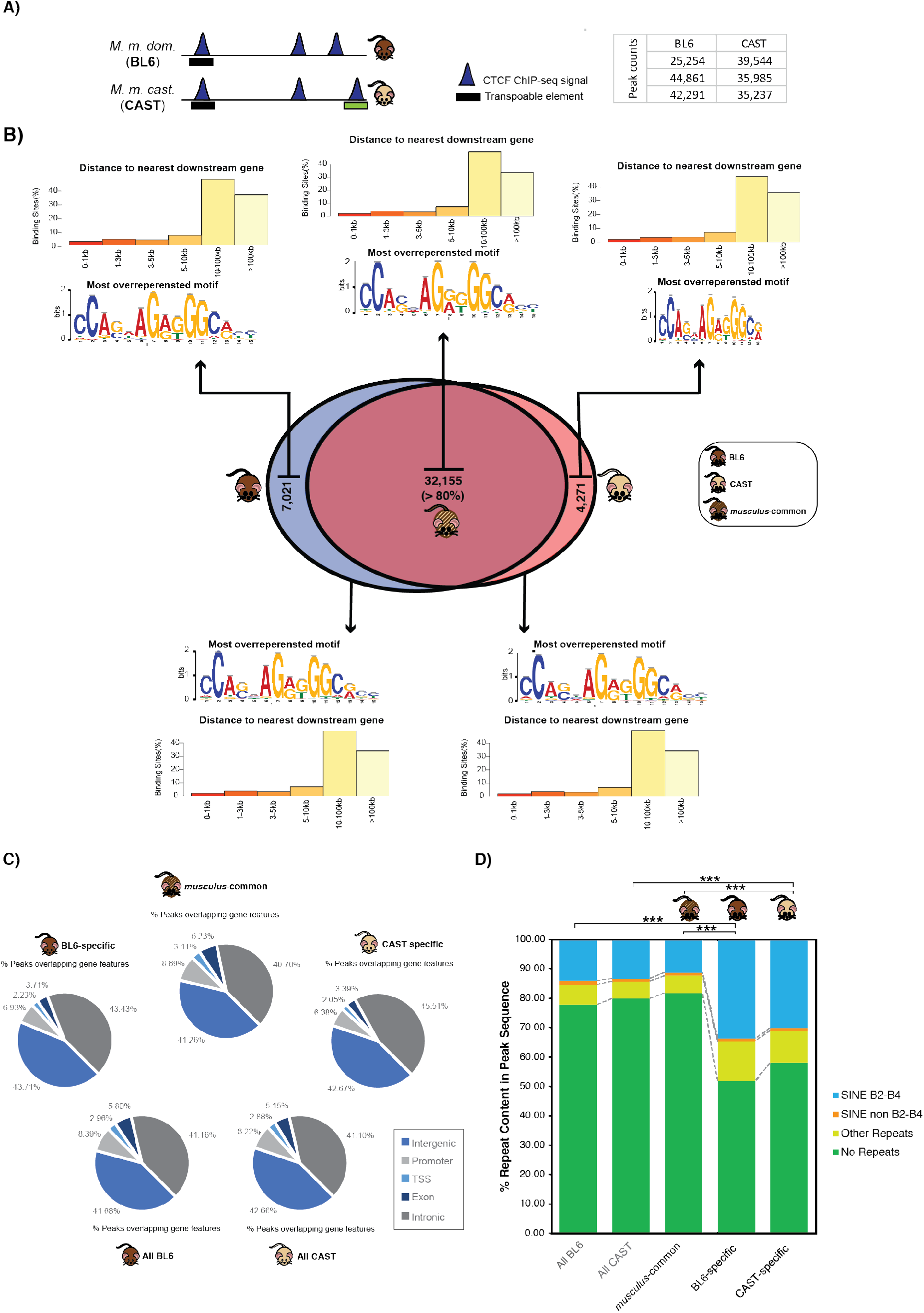
Overview of genomics features and evolutionary conservation of CTCF binding in the BL6 and CAST subspecies. **A)** schematic example of the contribution of transposable elements novel subspecies-specific CTCF binding. The peaks represent CTCF binding as determined from ChIP-seq experiments, while the boxes denote different groups of transposable elements (black = SINE, green = LTR). The table shows the peak counts (binding sites) retrieved from the three biological replicates for each subspecies. All downstream analysis utilised peaks common to a minimum of two replicates. **B)** The Venn diagram shows the degree of CTCF binding overlap in whole genome alignments between the *Mus musculus domesticus* (BL6) and *Mus musculus castaneus* (CAST) mouse subspecies. CTCF binding sites found aligned in orthologous locations are called *musculus*-common, while those with no alignment in the other species are subspecies-specific. For each evolutionary class of CTCF sites (above Venn diagram) and for all sites regardless of conservation between species (below Venn diagram), the most represented sequence motif and the distance to the nearest downstream genes. **C)** The pie charts show the gene features overlapping CTCF sites for all evolutionary classes in the Venn diagram in **B)**. **D)** The repeat content of all CTCF binding sites, and each evolutionary category described in B is measured as the percentage of a CTCF binding sites’ sequence that overlaps a repeat element. The asterisks indicate the significance of enrichment of SINE B2-B4 elements between subspecies-specific sets and all CTCF binding sites for both species and the *musculus*-common set (binomial tests, ***p < 0.0001)

## RESULTS

To study the evolution of CTCF binding between closely related species, we performed chromatin immunoprecipitation followed by high-throughput sequencing (ChIP-seq) in liver samples from two mouse subspecies: BL6 and CAST (Methods). We used three biological replicates for each subspecies and retained only those peaks present in at least two individuals, yielding comparable numbers of CTCF binding sites in both subspecies (Figure 1A). We performed evolutionary analysis of CTCF binding by finding those sites that occurred in orthologous locations in a pairwise alignment between the BL6 and CAST genomes. We found that the vast majority (>32,000) of CTCF binding occurs in orthologous locations, which we refer to as *musculus*-common sites (Figure 1B), in line with previous studies across more diverged mammals (29) and rodents (44). However, even within these closely related subspecies a considerable amount of biologically reproducible CTCF binding was found at subspecies-specific locations. In total, we identified in excess of 11,000 subspecies-specific CTCF binding sites in BL6 and CAST.

We next examined the genomic characteristics of subspecies-specific and *musculus*-common CTCF binding sites. Analysis of binding site positions relative to transcription start sites (TSSs) revealed an almost identical genomic location distribution, regardless of evolutionary conservation, with the largest portion of CTCF binding more than 10 kb from the nearest TSS (Figure 1B). We then classified each CTCF binding site by the genomic feature it overlaps (Figure 1B) and again found little differences between different categories of conservation. As expected, around 41-43% of CTCF binding within both *musculus*-common and subspecies-specific sites is intergenic, and the rest occurs within promoters or genes (45). Similarly, the canonical CTCF binding motif was retrieved from all CTCF binding sites (Figure 1C). Analyses of the genomic features of *musculus*-common and subspecies-specific CTCF binding suggests that the recently evolved sites perform similar functions to conserved sites.

### Transposable elements are responsible for half of subspecies-specific binding

Given the known contribution of transposable elements to CTCF binding site evolution (32, 35), we quantified the transposable element content in CTCF binding sites. Both subspecies had comparable overall repeat profiles with approximately 21% of all CTCF binding occurring within a repetitive element (Figure 1D), mostly of the B2-B4 rodent-specific subfamily of SINEs. However, for both BL6 and CAST, subspecies-specific binding sites were significantly more enriched in repetitive elements than *musculus*-common sites. Specifically, 34% of subspecies-specific CTCF binding overlapped a transposable element of the B2-B4 subfamily, more than a two-fold increase compared to all sites overlapping B2-B4 (14%) in both BL6-specific (BL6-specific vs. All BL6: binomial test p-value=3.12766E-07; BL6-specific vs. *musculus*-common: binomial test p-value=8.96592E-10) and CAST-specific binding sites (CAST-specific vs. All CAST: binomial test p-value=7.41176E-06; CAST-specific vs. musculus-common: binomial test p-value=1.85083E-07) (Figure 1D). The contribution of the SINE B2-B4 subfamily to subspecies-specific CTCF binding sites is likely an underestimation; an additional 15% of sites identified in only one biological replicate overlap with SINE B2-B4 and most likely represent weaker and/or CTCF binding variable between individuals.

To investigate how repeat element expansion contributed to the binding of other transcription factors expressed in the liver, we performed analogous evolutionary and repeat content analyses for the liver-specific transcription factors CEBPA and FOXA1. We first reanalysed publicly available ChIP-seq experiments (46) to identify *musculus*-common and subspecies-specific binding in BL6 and CAST as described above for CTCF (Methods). In both subspecies and for both liver-specific transcription factors, repeat elements contributed more to subspecific-specific than *musculus-*common binding, but to a much smaller extent than for CTCF (Figure 2A). Unlike CTCF binding, there was almost no overlap of CEBPA or FOXA1 with SINE B2-B4 elements (Figure 2B) or any other specific transposable element group. These results further confirm earlier studies that show SINE B2-B4 elements are specialised for the expansion of CTCF binding sites, and contribute to a larger portion of subspecies-specific binding than tissue-specific transcription factors.

**Figure 2.**
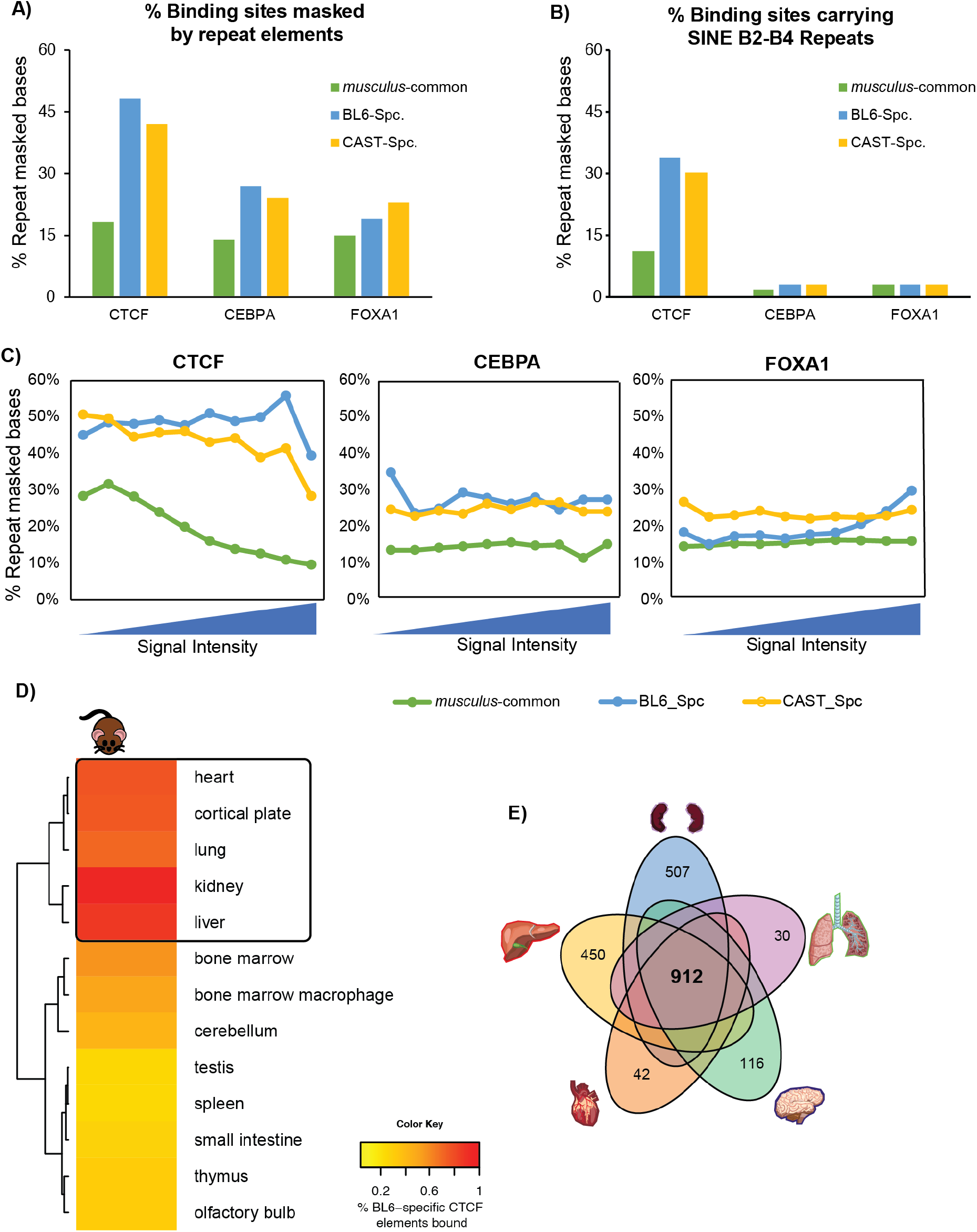
Repeat content and tissue distribution analysis of CTCF binding sites. **A)** Comparison of the total repeat element content in CTCF, CEBPA and FOXA1 transcription factor binding sites between sites with orthologous binding in the other subspecies (*musculus*-common) and subspecies-specific sites (BL6- and CAST-specific). All differences between subspecies-specific binding enrichment in repeat content with either the repeat content of all TF sites or the *musculus*-common set where found to be statistically significant (Binomial test, all p-values < 2.2e-16). **B)** Comparison of the total content of the SINE B2-B4 transposable element subfamily in CTCF, CEBPA and FOXA1 transcription factor binding sites between sites with orthologous binding in the other subspecies (*musculus*-common) and subspecies-specific sites (BL6- and CAST-specific). **C)** The dependence of repeat element content of binding sites on signal strength for CTCF, CEBPA and FOXA1 transcription factors. Within each transcription factor set, the data is binned in 10% bins based on binding site signal strength as estimated from the number of ChIP-seq reads mapped. **D)** Tissue distribution analysis of CTCF binding sites found to be subspecies-specific using ENCODE CTCF ChIP-seq data across 13 tissues. The heatmap indicates the overlap of BL6-specific CTCF binding with binding from ENCODE tissues, with the five most similar tissues highlighted. **E)** Venn diagram highlighting CTCF BL6-specific sites that are shared between all of the five most similar tissues and those found in only one of the tissues.

We next investigated the relationship between the age of transposable element insertion, as estimated by the repeat content within peaks, and transcription factor binding strength for all three transcription factors. We measured the repeat content of binding sites at increasing experimentally-determined ChIP-seq signal intensities (Methods) and found their genomic distribution to be indistinguishable from those CTCF binding sites with higher intensity (Figure 1B). In the *musculus*-common set of CTCF binding sites, the repeat content noticeably drops at increased binding intensities (Figure 2C) suggesting that older transposable elements have higher affinity for CTCF binding. In contrast, in the subspecies-specific sets, the overall repeat content and the SINE B2-B4 content were comparable across all ChIP-seq signal intensities (Figure 2C). The repeat content of the tissue-specific transcription factors CEBPA and FOXA1 was also independent from ChIP-seq signal intensity and similar across both *musculus*-common and subspecies-specific sites (Figure 2C). Interestingly, most tissue-specific transcription factor binding and *musculus-*common CTCF binding occurred in sites with noticeably smaller repeat content than subspecies-specific CTCF binding. This suggests that functional subspecies-specific CTCF binding sites have arisen from transposable elements more recently than tissue-specific transcription factors.

Taken together, these results demonstrate the extent and speed at which transposable elements can shape transcription factor binding. In just one million years of evolutionary time separating BL6 and CAST transposable elements have apparently contributed to almost half of the subspecies-specific CTCF binding profiles.

### Subspecies-specific CTCF binding is predominantly tissue-restricted

Although CTCF binding is more consistent across tissues than most other transcription factors, many binding sites are tissue-specific (47). To investigate the relationship between subspecies-specific binding and tissue-specificity, we determined the tissue distribution of binding sites across adult mouse tissues. We reanalysed ENCODE CTCF ChIP-seq data for BL6 adult male mice from 13 tissues: liver, lung, bone marrow, bone marrow macrophages, cortical plate, cerebellum, heart, kidney, thymus, spleen, olfactory bulb, small intestine and testis (47). We calculated the extent of CTCF binding overlap between ENCODE tissues and sites we found to be BL6-specific and show that a substantial subset are also bound in multiple other tissues (Figure 2D). Almost all BL6-specific binding sites we identified in liver are also present in the independent ENCODE liver experiments, showing that there is very little inter-individual variation for these sites. We selected the top five ENCODE tissues by the number of shared CTCF binding with our BL6-specific set for further analysis (Figure 2E). As expected, ENCODE liver has the most overlap with our liver datasets and kidney has the next most similar binding profile. For the BL6 binding sites we identified as *musculus*-common 67-85% are bound in these five tissues, compared to only 26-49% of the BL6-specific sites. The CTCF binding sites shared across at least one of the five tissues constitute 13% of all of BL6-specific CTCF sites, with only 4% (912) being bound in all tissues. Thus, these results show that subspecies-specific CTCF binding has relatively little variation between individuals and is more tissue-restricted than *musculus*-common sites.

### Subspecies-specific CTCF sites have genomic distribution similar to tissue-shared sites

To gain insight into the possible functional roles of evolutionarily distinct CTCF binding classes, we examined the genomic features near *musculus*-common CTCF sites and species-specific sites that were either tissue-restricted or tissue-shared. We first calculated the distance from each CTCF binding site to the transcriptions start site (TSS) of the nearest downstream gene. Regardless of the type evolutionary class or tissue-distribution of the binding site, we observed a large proportion of sites near the TSS (median −11 kb) (Figure 3A). The majority of the remaining sites lie further away from the TSS, more than 100 kb upstream. CTCF binding is depleted directly downstream of the TSS within the gene body.

**Figure 3.**
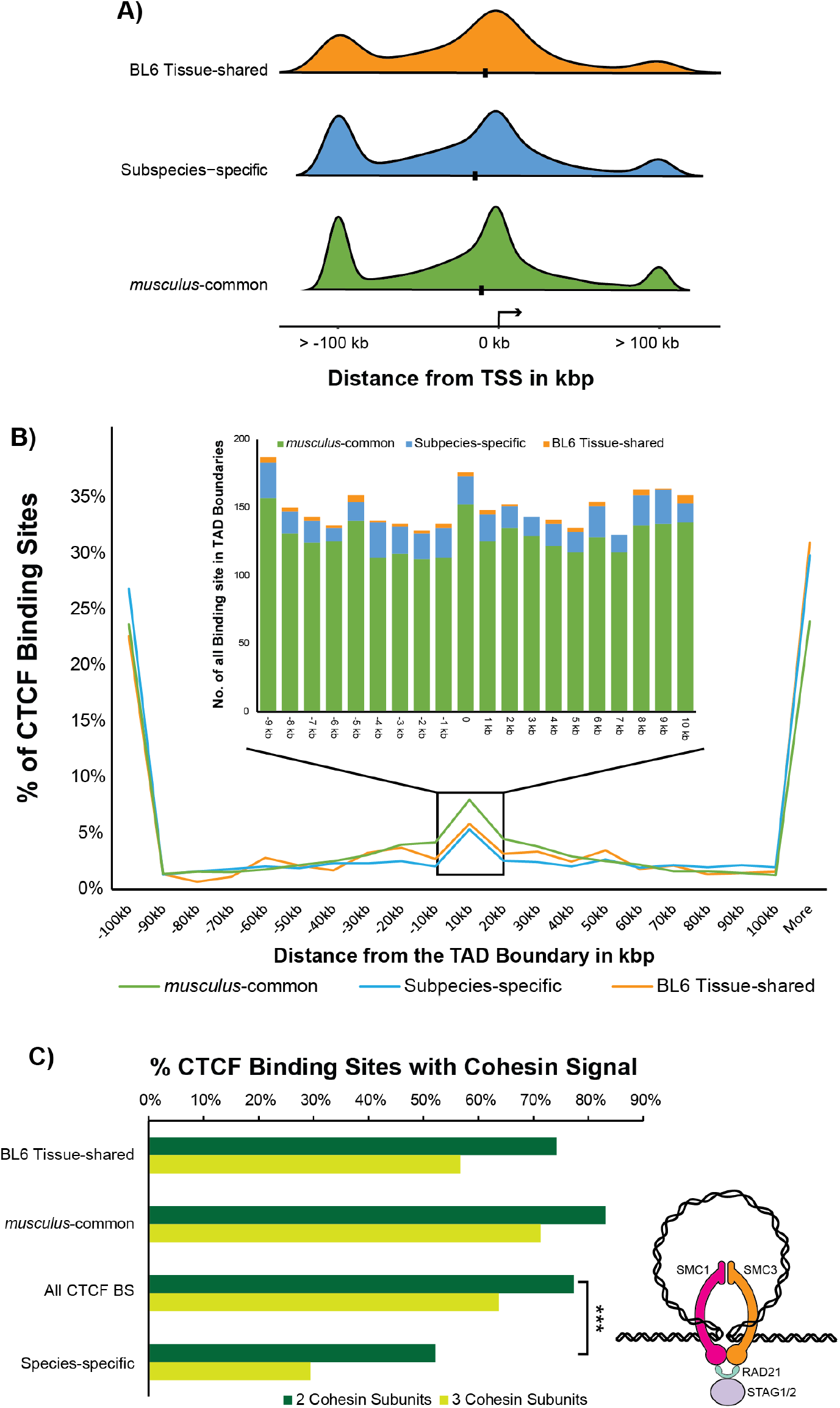
Characteristics of BL6-specific CTCF binding sites differ between tissue-shared and tissue-restricted sites. **A)** Distribution of the distances of CTCF binding sites to the transcription start site (TSS) of the nearest downstream gene based on their evolutionary class and tissue-distribution. The median is marked with a black point. **B)** Plot of the distances of CTCF binding sites to the nearest topologically associated domain (TAD) boundary reported in Vietri-Rudan *et al.* 2015 (29) for each evolutionary type of site. The inner box focuses on the region −10 kb and +10 kb from the nearest TAD boundary and shows the total number of CTCF sites. **C)** The percentage of all CTCF binding sites and different evolutionary classes of sites for which colocalisation with a cohesin complex protein was found. The asterisks indicate the significance of a Chi-square goodness of fit test for 2 cohesin subunits colocalising with CTCF between all CTCF sites and those found to be subspecies-specific (p-value = 2.8×10^−9^). The schematic diagram next to the bars is an overview of the structure of the cohesin complex.

We next examined the possible contribution of different evolutionary classes of CTCF to large-scale 3D genome structure. We took advantage of HiC experiments that determined the position of topologically-associated domain (TAD) boundaries in liver (29) to analyse the distribution of CTCF binding sites in relation to TADs. The distribution of CTCF binding distances to TAD boundaries is similar between all evolutionary categories of binding sites (Figure 3B). Specifically, the majority of both *musculus*-common and species-specific CTCF binding sites were located far (> 100 kb) from TAD boundaries, with slight enrichment around the boundaries. To explore binding at TAD boundaries in more detail, we limited our analysis to 10 kb around TAD boundaries. As expected, the enrichment of *musculus*-common sites at TAD boundaries was greater than for species-specific sites (29, 44). Though the majority of CTCF binding sites at TAD boundaries are *musculus*-common, subspecies-specific had similar enrichments at TAD boundaries (Figure 3A), regardless of their tissue-distribution. This suggests that some TAD boundaries, despite mostly being invariant across tissues (30), may be in part maintained by tissue-specific CTCF binding.

These observations illustrate that recently evolved, subspecies-specific CTCF binding sites mirror the pattern of *musculus*-common sites in their distribution around genomic features and may perform similar functions regardless of tissue-specificity.

### Recently evolved tissue-shared CTCF binding efficiently recruits cohesin

To form both large-scale and smaller-scale 3D genome structure, CTCF can help stabilise cohesin and form a chromatin loop (Figure 3C *diagram*). To quantify the extent to which subspecies-specific CTCF binding can recruit cohesin, we determined the level of co-location of CTCF and cohesin in BL6 mice. We used our previously-published ChIP-seq data from two biological replicates of adult mice livers for three proteins that form the cohesin complex: RAD21, STAG1 and STAG2 (48). RAD21 is necessary for the formation of the core ring of the cohesin complex, which is completed with either STAG1 or STAG2 (49, 50). We defined colocalised events as those where at least two cohesin subunits overlap with CTCF binding. All classes of CTCF binding sites colocalised with cohesin, with the highest fraction of colocalisation (~ 80%) observed for *musculus*-common CTCF sites. A significantly smaller portion of BL6-specific CTCF sites (~50%) colocalised with at least two cohesin subunits (Chi-square test p-value = 2.8×10^−9^). However, the tissue-shared subset of the BL6-specific sites (i.e. the ones bound in all five tissues in Figure 2C) colocalised with cohesin at the same level as the set of all CTCF binding sites, and only slightly less than that of the *musculus*-common sites (Figure 3C). The increased ability of BL6-specific tissue-shared binding sites to recruit cohesin in comparison to their more tissue-restricted counterparts suggest that there is a fundamental difference in the function of these recently evolved CTCF binding sites.

We investigated whether these BL6 tissue-shared CTCF-cohesin regions may participate in novel looping interactions and found no clear evidence. We speculated that BL6 tissue-shared CTCF-cohesin regions could act as novel loop anchors within established TADs and so limited our analysis to TADs with a minimum of two of these regions. There were fewer than 100 such regions in the whole genome and more than half of the potential novel loop anchors are found in single sites per TAD. We suggest that these sites are either stabilising existing TAD structures (44) or are forming intra-TAD domain loops that are below the resolution of the hi-C data available to us.

### CTCF contributes to a lineage-specific interferon gene and regulatory expansion

We next examined the genomic positions of CTCF binding sites colocalised with cohesin, and noticed an extreme but interesting example. Chromosome 4 of the BL6 genome contains a cluster of 15 CTCF-cohesin sites colocalised and tissue-shared binding sites within 58 kb, with all sites of similar lengths and with comparable ChIP-seq signal strength (Figure 4A). There is no detectable presence of either *musculus*-common or tissue-specific CTCF binding, indicating that this region is uniquely bound in a subspecies-specific and tissue-shared manner. This genomic cluster is unlikely to be the result of a genome assembly artefact as it is contained within a single clone within the reference BL6 mouse genome assembly (https://www.ebi.ac.uk/ena/data/view/AL928605). Interestingly, each of the 15 CTCF-cohesin sites within this region is upstream from a transcription start site of an immunity-related gene of the type 1 interferon zeta (Ifnζ) family. We have manually re-evaluated the annotation for all of the genes in the cluster and found the annotation to be largely correct (see Methods). The majority are novel/predicted protein coding genes and part of the comprehensive GENCODE annotation (51), with supporting transcript level evidence (Kawai *et al.* 2001; Okazaki *et al.* 2002). The 15 Ifnζ genes (also known as limitin) have previously been identified as a mouse-specific gene family expansion (52, 53).

**Figure 4.**
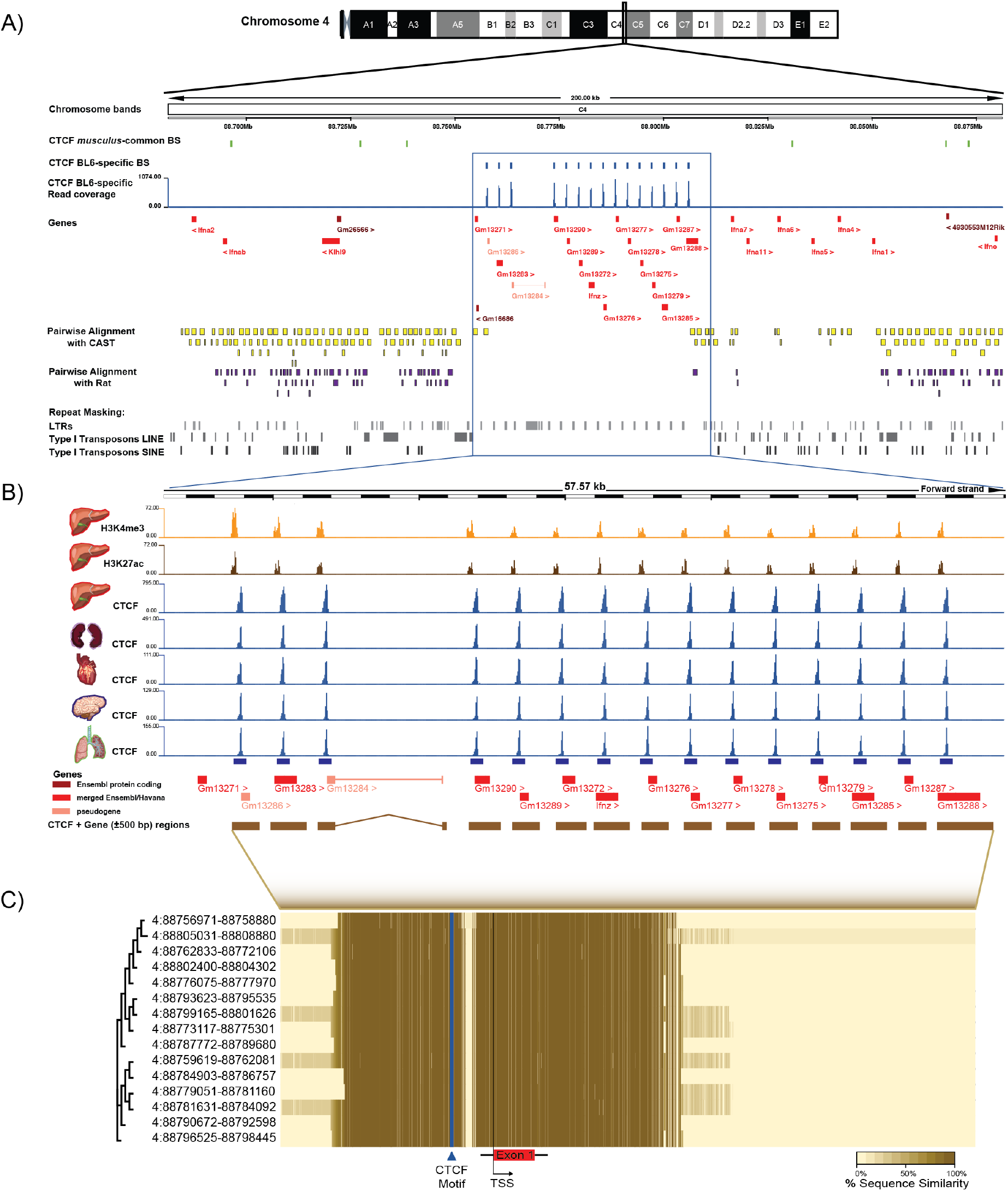
Evidence of a tandem duplication event of BL6-specific CTCF binding sites on Chromosome 4 in multiple tissues linked to the expansion of a family of interferon genes. **A)** A summary view of 200 kb on Chromosome 4 band C4 of the BL6 genome. The top two tracks show the CTCF-cohesin bound genomic regions in *musculus*-common and BL6-specific tissue-shared sites. The next track in blue shows the read coverage for CTCF BL6-specific tissue-shared binding sites. All genes in the 200 kb window are denoted below in red, with arrowheads indicating the direction of transcription. The pairwise alignments of the BL6 region to CAST (yellow) and rat (purple) show a noticeable lack of any orthologous regions in either species. The repeat content of the genomic region is shown in the bottom three grey tracks, illustrating the noticeable lack of any large scale repeat elements in the highlighted region. **B)** A zoomed-in view of the 57.6 kb region 4:88752534-88810107 in which BL6-specific CTCF-cohesin colocalised binding was observed. The top two tracks in orange and brown indicate read coverage signal from H3K4me3 and H3K27ac, respectively. The next five tracks in blue indicate CTCF ChIP-seq read coverage in five tissues (described in Figure 2) and illustrated that these CTCF sites are also bound in other tissues. The 15 interferon zeta cluster genes are shown below the tracks. **C)** Heatmap of the extent of sequence between similarity in a multiple sequence alignment of the 15 CTCF-cohesin binding sites and interferon zeta genes on the C4 band of BL6 chromosome 4. The dendrogram on the left is generated from the overall sequence similarity of each region. Coordinates represent the start and end positions of each binding site.

To more closely investigate the evolutionary origin of this gene cluster, we examined whole genome alignments within the rodent clade. Most of the BL6 gene cluster have no alignable regions in the genome of CAST, rat (Figure 4 A) or any of the other 13 mouse strains/species in pairwise alignments available in Ensembl version 91 (54). This cluster was characterised by a strikingly different distribution of transposable elements compared to the neighbouring regions. There was no detectable SINE or Long Interspersed Nuclear Elements (LINE) transposons, with the only repeat elements present belonging to the Long Terminal Repeat (LTR) ERVK subfamily (Figure 4 A). These LTR-ERVKs were between 450 and 550 bp in length and located in intergenic regions, around 500 bp up/downstream from gene bodies. The only detectable repeats in this region were six short simple repeats within (average length 50 bp), which are unlikely to be from transposable elements.

The LTR-ERVK elements did not colocalise with binding of the CTCF-cohesin complex or CTCF alone. The LTR-ERVKs, CTCF-cohesin bound regions and genes in this cluster exhibit high sequence similarity for large portions of their lengths (Figure 4 C). Given that LTRs have been shown to have regulatory activity in gene family expansions (55) we examined the enrichment of specific histone modifications from ENCODE (47). Each CTCF-cohesin binding site was also found to overlap in liver with H3K27ac predictive of regulatory activity (56) and H3K4me3 predictive of promoter function (57) (Figure 4 B). Examination of RNA-seq data for the thymus, spleen and liver available in Ensembl show transcriptional activity for all genes in this cluster, albeit at low levels (*data omitted from figure for space considerations*). These results show that the Ifnζ gene cluster is not only *musculus*-specific, but a more recent subspecies-specific tandem duplication event restricted to BL6. The gene cluster expansion and/or regulation of gene expression might have been facilitated by the presence of recently evolved transposable elements.

Interestingly, despite the lack of alignable regions in more closely related species, LASTz whole genome alignments with other eutherian mammals revealed that 14 of the 15 genes in the BL6-specific cluster are aligned with high coverage (50-100%) to a single gene in the pig (54, 58). The predicted pig protein-coding gene is located on chromosome 1 (ENSSSCG00000039987) and has 13 paralogues within a 470 kb cluster, albeit separated by 24 intervening genes (Supplementary Figure S1). All the genes belong to the Ensembl protein family PTHR11691 (Interferon Precursor), which has only a single member outside the cluster. Similarly to the BL6 cluster, this region is enriched with LTRs punctuating the intergenic regions. Compared to the surrounding genomic regions, this pig cluster has lower GERP conservation scores with less constrained elements, indicating a more recent evolutionary origin. Motif discovery analysis of 1kb regions around the 14 gene paralogues revealed that almost all have a CTCF binding motif less than 100 bp from the transcription start site (Figure 5C). These results support previous suggestions of lineage-specific expansions of IFNδ in the porcine lineage and IFNζ in the mouse lineage from a more ancient IFN gene (53) and may be an example of convergent evolution.

## DISCUSSION

Recent studies have demonstrated the regulatory potential of species-specific transposable element insertion in primates, especially in tissue-specific contexts (59, 60). In particular, the emergence of novel CTCF binding by repeat expansion is a mechanism known to have repeatedly reshaped the genomes of highly diverged mammals (32, 35). In the mouse genome, very recent waves of expansions of SINE B1 and SINE B2 transposable element subfamilies are known to have created many novel CTCF binding sites not present in rat (37, 61). Here, we use two closely related mouse subspecies, *Mus musculus domesticus* (BL6) and *Mus musculus casteanus* (CAST), separated by only 1 million years of evolution to reveal the speed of repeat expansion associated CTCF binding and to suggest potential functions for these young sites.

It has previously been shown that a large fraction of hominidae-specific transcription factor binding sites, when compared to ancestral human-mouse shared ones, are enriched near genes implicated in specific pathways and may therefore have distinct biological functions (62). However, mouse and rat species-specific CTCF sites have comparable functional effects to shared sites on chromatin domain demarcation and transcriptional regulation (35). To investigate the potential biological function of even younger transcription factor binding sites, we compared the genomic locations and gene function enrichment of subspecies-specific CTCF sites versus the sites common between BL6 and CAST. We found that they are mostly indistinguishable in the gene features they bind, distance to transcription start sites or TAD boundaries. Our results illustrate how these evolutionarily young CTCF sites have been captured into operational regions of the genome and adopted functions similar to *musculus*-common sites. This observation is in line with the observation that species-specific CTCF binding sites cluster with species-shared sites to provide functional redundancy (44). The final conclusions on the biological function of lineage-specific CTCF sites will require more targeted *in vivo* studies, such as CRISPR-Cas9 of individual CTCF sites or conditional knockdowns of CTCF.

It has also been shown that tissue-shared CTCF binding is more conserved than tissue-specific CTCF binding (62, 63), but the functional differences between tissue-shared and tissue-specific young CTCF sites have not yet been investigated. By determining the tissue-distribution of CTCF binding of evolutionarily young CTCF sites, we found significant differences between tissue-shared and tissue-restricted sites. Most subspecies-specific sites are restricted in their tissue distribution, but many are still bound across multiple tissues originating from all three germ layers. These subspecies-specific tissue-shared sites are almost as likely to be colocalised with cohesin as *musculus*-common, and far more than other subspecies-specific sites. This suggests that these tissue-shared, subspecies-specific sites have a greater regulatory potential and are more likely to adopt functional signature than their cell-type specific counterparts. Due to the resolution of published Hi-C experiments for determining 3D genome structure, we could not establish whether pairs of colocalised CTCF-cohesin subspecies-specific sites are implicated in forming chromatin loops. Most pairs of sites were too close to *musculus*-common CTCF-cohesin regions, or too close to each other, to be able to establish their contacts. Chromatin capture experiments with higher resolution would be needed to investigate loops associated with these sites and to establish the extent to which CTCF subspecies-specific sites contribute to transcriptional regulation either in tissue-shared or tissue-restricted scenarios.

There have been previous reports of clustered expansion of functional genes of the interferon alpha family between the BL6 and 129/5v mouse strains (64). Similarly, the expansion of the *Abp* gene cluster is well described in mice and is associated with transposable elements (37, 55). We found an example of a BL6-specific gene cluster and CTCF binding expansion of the type 1 interferon zeta family is associated with a specific LTR expansion, and evidence of a similar expansion in pig. The LTRs may either have served to duplicate the genes through non-allelic homologous recombination or helped provide binding sites for regulatory factors (65). Within this gene cluster, subspecies-specific CTCF binding colocalised with cohesin and histone modifications indicative of active promoter function, and the genes show transcriptional activity. This suggests that CTCF may have helped established 3D genome structure and transcriptional regulation within the locus. The mouse gene cluster has not been well-described in the literature, though it is known to be a mouse-specific expansion (53). Characterisation of the larger area around the subspecies-specific gene cluster revealed that this interferon locus in the mouse has a number of orthologues in the human interferon locus on chromosome 9, but the smaller subspecies-specific cluster was completely overlooked in part due to concerns of an assembly artefact (64, 66). Our detailed manual curation of the genes within the cluster, and confirmation that the region is contained within a single BAC clone, disprove assembly artefacts in the region. Further exploration of the genes in the locus is difficult due to many subsequent changes to names or gene IDs, and due to changes to the protein-coding status of some genes as transcriptional evidence improved.

Our results demonstrate that evolutionarily young CTCF binding sites establish functional signatures and how these elements contribute to genome regulation including at lineage-specific gene clusters.

## METHODS

### Data Sources

Liver ChIP-seq libraries from two closely-related *Mus* subspecies were obtained for *Mus musculus domesticus* (C57BL/6J *or* BL6) from Thybert et al 2018 (37), and for *Mus musculus castaneus* (CAST) from Kentepozidou et al 2020 (44), each with three biological replicates. Liver ChIP-seq libraries for CEBPA and FOXA1 were retrieved from Stefflova et al. 2013 (46) for both subspecies in biological triplicates. ChIP-seq data for three cohesin-complex subunits (Rad21, STAG1 and STAG2) in liver, two biological replicates for each subunit, from adult male mice and matched controls were retrieved for BL6 from Faure et al. 2012 (48). TAD boundary domains were retrieved from Vietri Rudan *et al.* 2015 (29). CTCF and histone modifications H3K4me3 and H3K27ac ChIP-seq libraries across adult mouse tissues were retrieved from the ENCODE Project data repository for BL6 adult (8 weeks) male mice and 13 tissues: liver, lung, bone marrow, bone marrow macrophages, cortical plate, cerebellum, heart, kidney, thymus, spleen, olfactory bulb, small intestine and testis (47).

### ChIP-seq Sequence Alignment and Peak Calling

All libraries were retrieved as raw ChIP-seq FASTQ reads were subject to quality control using standard parameters in FastQC version 0.11.5 (67). Good quality reads (min Phred score >= 30) were subsequently aligned to most recently available genome assembly in Ensembl (GRCm38 for BL6 and CAST_EiJ de novo assembly downloaded from ftp://ftp-mouse.sanger.ac.uk/, and available in Ensembl version 84 for CAST). We aligned the sequence reads to the reference genomes using BWA version 0.7.12 (68) for each biological replicate and control. Aligned reads were afterwards filtered for duplicate and non-unique reads, sorted and indexed using SAMtools version 1.2 (69). CTCF binding sites were identified by peak calling from aligned sequence reads using MACS version 2.1.0 (70) with a p-value threshold of 0.001 to call peaks representing CTCF bound regions. Peaks found in at least two biological replicates out were used for downstream analysis. Motif analysis focused on the summit point (±50 bp) of each identified CTCF binding sites using the MEME suite version 4.10.2 (71, 72). The most overrepresented motif found in each dataset is reported in Figure 1B. CTCF binding sites characterisation in terms of gene feature occupancy and proximity to downstream gene bodies was performed using the annotatePeaks.pl tool from HOMER (Hypergeometric Optimization of Motif EnRichment) suite (v4.11) with annotation from the most recent mouse genome assembly (GRCm38) (73).

### Interspecies comparisons

Interspecies comparison between BL6 and CAST was performed first using a multiple alignment of 15 de novo assemblies of laboratory and wild-derived strains genomes within *Mus musculus* (42, 74). Orthologous regions with a CTCF binding site present in orthologues alignment regions in both species was considered a “*musculus*-common” site, whilst sites found in only one of the species, but absent from the other, was considered “subspecies-specific”.

### Repeat Masking of CTCF binding sites

CTCF binding regions from *musculus*-common and species-specific sets of the data for both species were screened for repeat elements using RepeatMasker 4.0.5 (75) using the rodent repeat libraries from RepBase (v20140131) for the two murine species, with the cross_match search engine, masking for interspersed and simple repeats. Fragmented hits found to be part of the same repeat were merged as one.

For the repeat content analysis, all dataset for each transcription factor in both species were divided into ten 10% bins based on descending signal intensity of the ChIP-seq signal, and in each bin, repeat masking followed to determine the repeat content of each.

### Cross-Tissue Analysis of Species-Specific CTCF Binding

CTCF peaks overlapping between *musculus*-common/BL6-specific sites from our CTCF liver binding sites and ENCODE CTCF tissues’ binding sites were identified using BEDTools version 2.25.0 (76, 77). Tissue-shared BL6-specific CTCF binding sites were identified as the intersection of all BL6-specific binding sites from the top five ranking tissue above.

### Genomic Distribution Analysis

To calculate the distance from CTCF binding sites to the nearest up/downstream topologically-associated domain (TAD) boundary, liver TAD boundary data from Vietri Rudan *et al.* 2015 (29) were used, and distance determined using BEDTools (76, 77). CTCF binding sites regions were analysed with GREAT version 3.0 (78) using default parameters to determine the distance from each CTCF site of each category to the nearest transcription start site (TSS). All CTCF sites more than ±100 kb from the nearest TSS were pooled together.

### CTCF-Cohesin Colocalisation Analysis

Genomic regions where at least two cohesin subunits peaks overlap were merged using BEDOPS version 2.4.30 (79), and cohesin merged regions overlapping with *musculus*-common/BL6-specific/BL6-tissue-shared from our CTCF liver binding sites were identified. The intersection analysis was done for CTCF co-occupancy with two and three subunits, owing to the significantly fewer number of ChIP-seq peaks retrieved from the STAG1 data.

### Chromosome 4 Interferon-zeta gene-cluster Analysis

CTCF binding sites coordinates in *bed* format along with ChIP-seq coverage reads in those regions were uploaded for display on the Ensembl genome browser version 89 (54). These included reads from liver and the other four tissues, plus ChIP coverage reads from two histone marks for liver, H3K4ac27 and H3K4me3, from the ENCODE data repository. Ensembl genome browser was used to display gene annotations, pairwise alignments with CAST and Rat, repeat elements enrichments for transposons and LTRs and genomic annotations. Sequence similarity for the 15-gene cluster, upstream CTCF binding regions, and the complete 15 constructs of CTCF binding sites plus the gene sequence plus ±500 bp were determined using Clustal Omega (80), using default parameters.

We utilised the Comparative Genomics tool of the Ensembl Genome Browser to look at the BLASTz/LASTz whole genome alignment between the Chromosome 4 Interferon-zeta gene-cluster and all available pairwise alignments with other organisms (54). An orthologous gene was found in the pig genome whose target sequence matched 14/15 from the mouse cluster with Query %id of > 50%. We used BLAST/BLAT to scan the pig genome for paralogues to the gene based on sequence similarity. As with the mouse cluster, Ensembl genome browser was used to display gene annotations, repeat elements enrichments for transposons and LTRs and GERP scores. Next, we scanned the 1 kb sequences upstream of each gene's TSS for the enrichment of motifs using MEME (4.12.0), setting 5 as a maximum number of motifs, and a motif width between 6-50 bp. The top 5 motifs from all upstream sequences were subsequently submitted to TOMTOM (2.14.0) to search available databases for annotated motifs to match (71, 72).

The manual annotation review of the locus on chromosome 4 determined that the annotation of the region was essentially correct with only a couple of minor issues identified and corrected. Specifically, Gm16686 was identified as a spurious protein-coding gene that had already been removed from RefSeq and was flagged for removal in GENCODE release M19. RP23-400P11.4 was added as novel interferon pseudogene located at BL6 chr4:88754471-88754678 due to the clear pseudogenic characteristic of a significantly truncated 3' end. Finally, we reviewed Gm13286, which is annotated as a pseudogene in GENCODE, but considered protein-coding by RefSeq. It has a premature STOP compared to other family members, though it only loses the last 3aa of the typical protein. Based on the GENCODE annotation guidelines, Gm13286 is correctly annotated as a pseudogene although the coding status should be further investigated to make a definitive determination.

## ACKNOWLEDGEMENTS

We thank Maša Roller for support and for a critical reading of the manuscript and suggestions for its improvement; and Christine Feig for supervision and early discussions shaping the direction of the work.

## AUTHOR CONTRIBUTIONS

DA, DTO, and PF conceived and designed the study. DA led and conducted all analysis. JMM did the manual genome annotation. DA wrote the manuscript and created the figures, which were both refined and edited by DTO and PF. All authors read the final manuscript and provided critical comments.

## FUNDING

Funding is provided by the Wellcome Trust (WT202878/B/16/Z, WT108749/Z/15/Z, WT202878/Z/16/Z), Cancer Research UK (20412), the European Research Council (615584), the National Human Genome Research Institute of the National Institutes of Health (U41HG007234), the European Molecular Biology Laboratory, and the EMBL International PhD Programme.

## AVAILABILITY OF DATA AND MATERIALS

All mouse ChIP-seq libraries utilised in this study are readily available through the Array Express repository (https://www.ebi.ac.uk/arrayexpress/) under these accession numbers: E-MTAB-5769 (for BL6) (37), E-MTAB-8014 (for CAST) (44), E-MTAB-941 (Cohesin) (48), E-MTAB-1414 (FOXA1 and CEBPA) (46). The TAD data used in the analysis were obtained from Vietri Rudan *et al.* 2015 (29). ENCODE data was used for CTCF binding across tissues (47).

## ETHICS APPROVAL AND CONSENT TO PARTICIPATE

All animal procedures were conducted in accordance with the project (70/ 7535) and personal licenses, revised by the Animal Welfare and Ethical Review Body at Cancer Research UK Cambridge Institute and issued under the United Kingdom Animals (Scientific Procedures) Act, 1986.

## COMPETING INTERESTS

PF is a member of the Scientific Advisory Boards of Fabric Genomics, Inc., and Eagle Genomics, Ltd. All other authors declare that they have no competing interests.

---

**Supplementary Figure 1.**
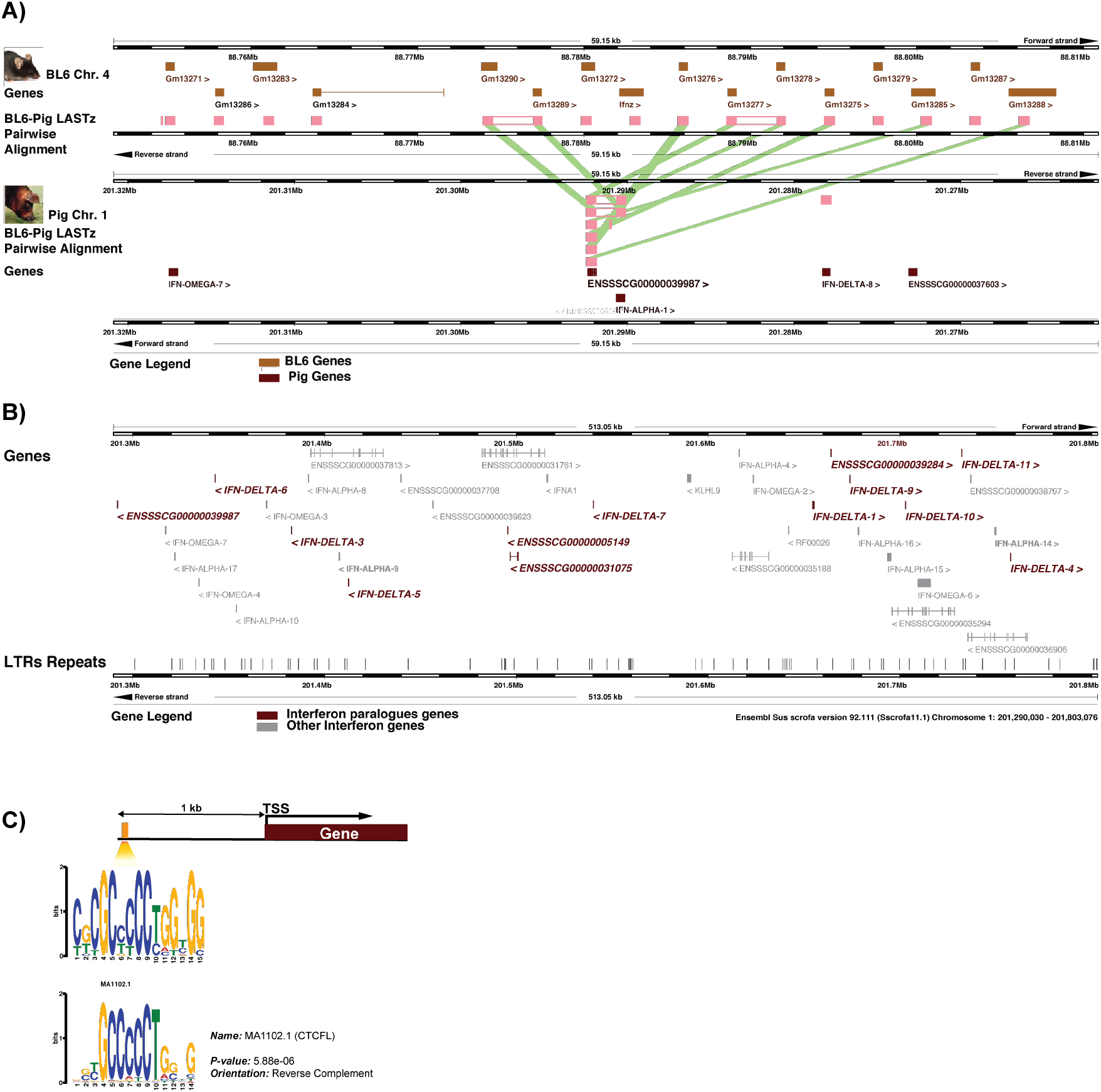
Convergent evolution of an orthologous interferon gene cluster in pig. **A)** Genome browser display of BL6-Pig LASTz pairwise alignment of the 15-gene cluster. Pink tracks show the BL6 genome regions aligning to sequences in the pig genome. **B)** A zoom-in view of the orthologous gene cluster of interferon precursors in the pig genome. The orthologous gene in (A) is shown as the leftmost gene in the window in brown italics. The 12 paralogues to this gene are highlighted in brown italics with the other interferon genes in light grey. The arrowheads indicate the direction of transcription. The dark grey tracks at the bottom indicate the LTR repeat content of the region. **C)** A schematic diagram showing the position of the CTCF motif enriched at around 1 kb from the TSS of 12/13 genes in the cluster, with the motif composition below the orange track indicating the position. The motif underneath is the CTCF canonical motif with the p-value of the probability that the match occurred by random chance.

